# A Chemical Toolbox for the Study of Bromodomains and Epigenetic Signaling

**DOI:** 10.1101/391870

**Authors:** Qin Wu, David Heidenreich, Stanley Zhou, Suzanne Ackloo, Genevieve Deblois, Shili Duan, Kiran Nakka, Jeffrey Dilworth, Mathieu Lupien, Paul E. Brennan, Cheryl H. Arrowsmith, Susanne Müller, Oleg Fedorov, Panagis Filippakopoulos, Stefan Knapp

**Affiliations:** 1nstitute of Pharmaceutical Chemistry, Goethe-University Frankfurt, 60438 Frankfurt, Germany; Structural Genomics Consortium, University of Toronto, Toronto, Ontario, M5G 1L7, Canada; Princess Margaret Cancer Centre, University Health Network, Toronto, Ontario, Canada; Department of Medical Biophysics, University of Toronto, Toronto M5G 2M9, ON, Canada; Structural Genomics Consortium, Buchmann Institute for Life Sciences, Goethe-University Frankfurt, 60438 Frankfurt, Germany; German Cancer Network (DKTK), Frankfurt/Mainz, 60438 Frankfurt, Germany; Target Discovery Institute and Structural Genomics Consortium, University of Oxford, Oxford OX3 7DQ, UK; Sprott Centre for Stem Cell Research, Regenerative Medicine Program, Ottawa Hospital Research Institute, Ottawa, Canada

## Abstract

Bromodomains (BRDs) are evolutionary conserved epigenetic protein interaction modules which recognize (“read”) acetyl-lysine, however their role(s) in regulating cellular states and their potential as targets for the development of targeted treatment strategies is poorly understood. Here we present a set of 25 chemical probes, selective tool small molecule inhibitors, covering 29 human bromodomain targets. We comprehensively evaluate the selectivity of this probe-set using BROMOscan^®^ and demonstrate the utility of the set using studies of muscle cell differentiation and triple negative breast cancer (TNBC). We identified cross talk between histone acetylation and the glycolytic pathway resulting in a vulnerability of TNBC cell lines to inhibition of BRPF2/3 BRDs under conditions of glucose deprivation or GLUT1 inhibition. This chemical probe set will serve as a resource for future applications in the discovery of new physiological roles of bromodomain proteins in normal and disease states, and as a toolset for bromodomain target validation.

## Introduction

Genetic and epigenetic variation, as well as environmental and lifestyle factors, work in concert to influence human health and disease. In recent years the essential role of epigenetic modifications in regulating gene expression and cellular differentiation has emerged^1, 2^. Apart from changes in DNA methylation, covalent post translational modifications (PTMs) of histones and other nuclear proteins define a complex language, the epigenetic code, which regulates chromatin structure and dynamics. Lysine acetylation (Kac) is a major epigenetic PTM occurring on histone proteins, which has been studied broadly. Kac has generally been associated with activation of transcription through opening of chromatin structure, although some recent studies have found some Kac marks to be responsible for the compaction of chromatin, protein stability, and the regulation of protein-protein interactions^3^.

Disruption of histone acetylation patterns has been linked to the development of disease, which may occur through mutations that deregulate enzymes responsible for adding or removing these histone acetyl marks, as well as the protein interaction modules that recognize and interpret this important PTM^4, 5^.

Histone acetylation is a highly dynamic process that is regulated by histone acetyltransferase (HAT) and histone deacetylases (HDACs) that respectively “write” and “erase” acetylation marks. The complex pattern of acetylation marks is interpreted (“read”) by reader domains of the bromodomain (BRD) family of proteins. BRD-containing proteins are evolutionary conserved and of substantial biological interest, as components of transcription factor and chromatin modifying complexes and determinants of epigenetic memory. There are 61 BRDs expressed in the human proteome, present in 46 diverse proteins. However, some atypical bromodomains, which lack essential residues have little or no-activity towards Kac-containing histone sequences and may recognize other epigenetic marks or unmodified peptide sequences, while canonical BRDs may also bind to complex patterns of modification around a central Kac site that often contain other post translational modifications^6, 7^.

The modular nature of many BRD-containing proteins, which typically harbour a number of diverse reader domains in addition to enzymatic functionalities and role(s) as scaffolds in large chromatin modifying complexes, makes their functional study a challenging task. However, the development of highly selective inhibitors has provided versatile tools for functional studies on endogenous BRD-containing proteins which can now be used to unravel the role of the epigenetic Kac-dependent reading process in chromatin biology as well as in the development of disease. This is exemplified by the development of highly potent inhibitors for BET (Bromo and extra-terminal; BRD2, BRD3, BRD4, BRDT) BRDs^8, 9^, which has led to numerous translational and functional studies on this subfamily of bromodomain proteins^10–18^.

Chemical probes, small molecule tool inhibitors with selectivity against similar proteins, have led to the validation of many disease targets, making seminal contributions to our understanding of complex cellular processes. However, chemical probes need to be highly selective, cell active and therefore need to be comprehensively characterized in order to link observed phenotypic responses to targeted proteins. Unfortunately, selectivity and potency of tool compounds are often insufficient resulting in contradictory and erroneous results^19–22^. Following the disclosure of potent BET BRD inhibitors, other members of the BRD-family of interaction modules have been found to be highly druggable^23^, resulting in the identification of chemical fragments that were subsequently developed into potent and selective chemical probes^24–31^. However, BRDs outside the BET family have not been found to be major regulators of primary transcription control, posing challenges for the discovery of functional roles of these conserved domains^32^. As a result, only a few studies have reported phenotypic consequences inhibiting non-BET BRDs pointing to important roles in cellular differentiation^33, 34^.

Here, we characterized a comprehensive set of BRD chemical probes covering all subfamilies previously identified with good druggability scores^23^. Using a standardized commercial assay format (BROMOscan^®^), based on a high throughput binding assay originally developed to assess the selectivity of kinase inhibitors^35^, we evaluated the selectivity of this BRD chemical probe-set and determined a total of 626 K_D_ values on all detected interactions. We present here an overview of the binding modes of these inhibitors resulting in the excellent selectivity of these chemical probes. To exemplify the use of this probe-set in biological systems, we further screened this probe collection on a cellular model of muscle differentiation identifying BET BRDs as major regulators in this context.

Furthermore, systematic investigations of all existing BRD inhibitors in triple negative breast cancer (TNBC) cell lines have revealed an essential role of BRD inhibitors to target the metabolic vulnerability of TNBC, demonstrating their utility as a collection to uncover previously unknown crosstalk between epigenetic regulators and cell metabolism. Together, our study provides a comprehensive structural and functional insight on BRD inhibitors, establishing a powerful resource for future mechanistic studies of this family of epigenetic reader domains, and underscoring the broad utility and immediate therapeutic potential of direct-acting inhibitors of human bromodomain proteins.

## Results

### A set of highly selective bromodomain probes

BRDs have been grouped into eight major families based on sequence and structural homology^6^ (**Figure 1**). Potent chemical probes have now been developed for most of these families with the notable exception for BRDs of the PML (promyelocytic leukemia)-SP100 nuclear bodies (family V), which harbor a PHD-BRD tandem reader cassette. However, some families are still insufficiently covered, including members of families VI and VII which also have atypical and shallow Kac-binding pockets. In contrast, family I which contains the histone acetyltransferases PCAF and GCN5 as well as CECR2 and FALZ is well covered by chemical probes. The dual PCAF/GCN5 chemical probe L-Moses showed good potency for these two highly related bromodomains (K_D_ of 126 nM and 600 nM respectively, determined by ITC)^36^. GSK4027 offers an alternative chemotype to antagonize the BRDs in these two targets with improved potency (K_D_ 1,4 nM determined by BROMOscan^®^ for both BRDs)^37^. Early lead molecules for bromodomains of CECR2 and FALZ were discovered by screening a series of triazolophthalazines^38^. However, compounds of this series inhibited several BRD family members and exhibited poor solubility, limiting further development. NVS-CECR2-1 was the first potent chemical probe targeting CECR2 with good potency (80 nM, determined by ITC) and selectivity. An alternative probe molecule, GNE-886, has recently been published showing however some activity towards the BRDs of BRD9, BRD7 and TAF1/TAF1L^39^.

**Figure 1:**
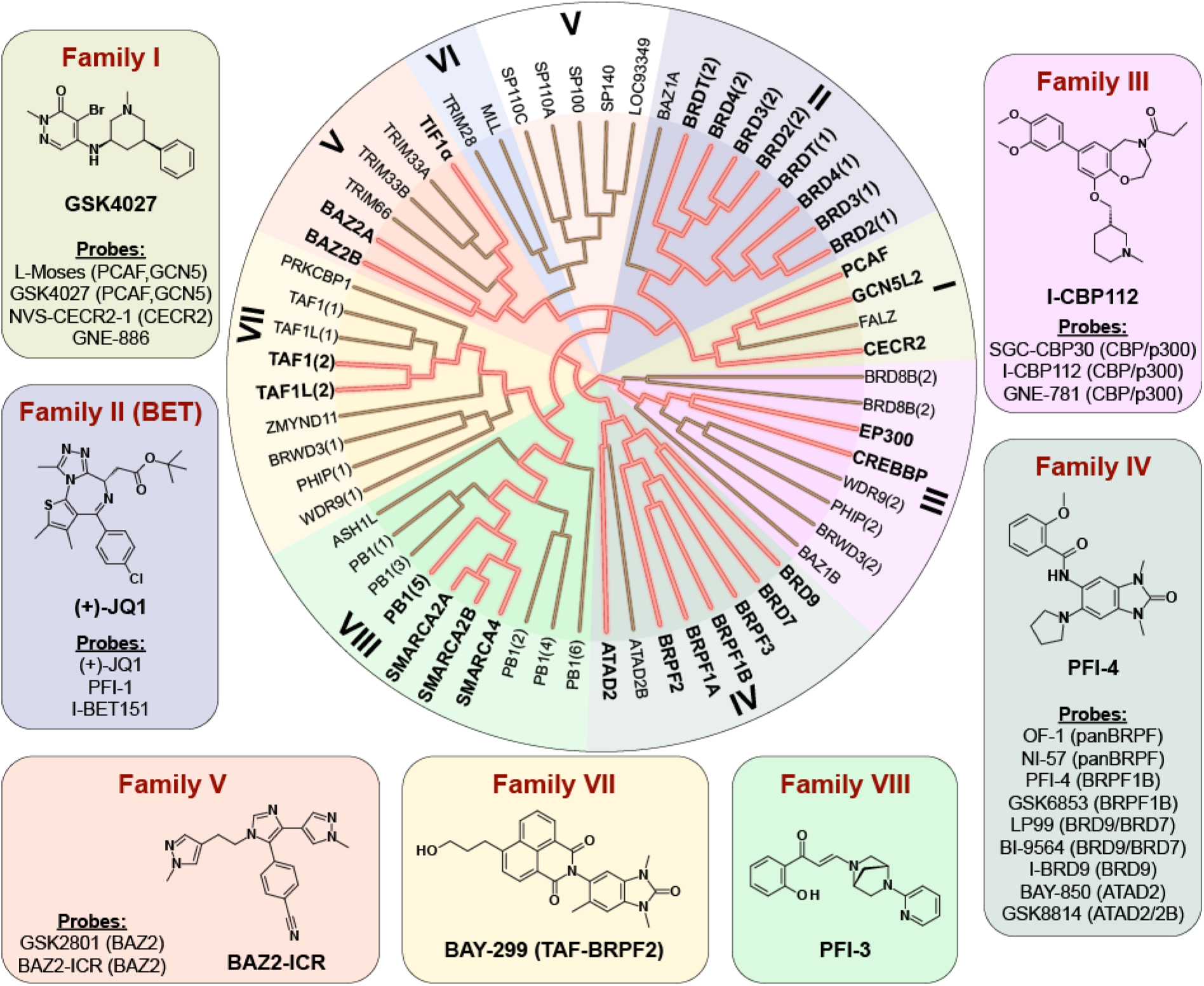
Chemical probes of the human bromodomain family. The set includes probes developed by our laboratory and a selection of additional inhibitors that are available. For each BRD family one example of a structure of chemical probe is shown. Additional probes are listed and a summary showing all chemical structures is included in Supplemental Table S1. BRD family members for which probes have been developed are highlighted in bold and by dark red lines in the dendrogram.

To date, the BET BRDs (Family II) have had the greatest activity in inhibitor development, undoubtedly due to the strong and clinically relevant phenotypes observed for these compounds. This is an area that has rapidly evolved and has been previously reviewed in detail^10, 40^. The first published Kac competitive BRD inhibitors which now have been widely used are the thienodiazepine (+)-JQ1, (henceforth, JQ1)^8^ the related clinical compound OTX015, as well as the benzodiazepine iBET^41^. Inhibitors of this family show panBET activity primarily against the first BRD with slightly lower binding affinity towards the second BRD in BET proteins. More recently, antagonists featuring diverse Kac mimetics have been developed, including the isoxazole I-BET151 (GSK1210151A)^42, 43^, the tetrahydroquinazoline PFI-1^44^ and the tetrahydroquinoline I-BET726 (GSK1324726A)^45^. Here we included in our probe set JQ1, I-BET151 and PFI-1 as three structurally diverse and unencumbered chemical probes for BET proteins.

Family III contains BRDs present in the histone acetyl-transferases (HATs) p300 and CBP, as well as a number of diverse BRDs for which no potent inhibitors have been identified so far. The first inhibitor developed for CBP/p300, SGC-CBP30, exhibited potent activity for BRDs in these two HATs (K_D_: 21 and 38 nM, respectively), retaining however significant BET activity, which needs to be taken into account in cellular assays by using appropriate concentrations^29, 46^. An alternative chemical probe is the benzoxazepine I-CBP112^34^. Recently, a highly potent antagonist, GNE-781, which has 650-fold selectivity over BRD4 for CBP/p300 became available^47^.

Family IV contains BRDs participating in HAT scaffolding (BRPF1-3), and chromatin remodelling complexes (BRD7, BRD9 and ATAD2A/B). Several chemical probes target BRD7 and BRD9, including BI-9564^48^, LP-99^26^, as well as I-BRD9 which has good selectivity towards BRD9 over BRD7^49^. The ATAD2 and ATAD2B BRDs have been targeted using the potent and selective antagonist GSK8814^50^, while the allosteric BRD antagonist BAY-850 was developed as a specific probe against ATAD2^51^. BRPF BRD antagonists are well represented by the pan-BRPF chemical probes OF-1 and NI-57, and the two related BRPF1B selective chemical probes PFI-4 and GSK6853^52, 53^. In addition, BAY-299, a dual activity chemical probe for BRPF2 and TAF1(2), has also been developed providing the only currently available chemical tool for family VII^54^. Similarly, compound 34, a dual activity antagonist of TRIM24 and BRPF1B represents the only chemical tool currently developed for TRIM24^55^.

Family VI is divided into the RING-Type E3 Ubiquitin Transferase of the TRIM (Tripartite Motif-Containing Protein) family and BAZ2 (Bromodomain Adjacent To Zinc Finger Domain) which are components of chromatin remodeling complexes. As mentioned above the dual chemical probe for the TRIM24 has recently been developed offering proof of principle that BRDs within this family are also druggable^55^. Two high affinity probes have also been developed against the BRDs of BAZ2A/B, GSK2801 and BAZ2-ICR^25, 27^. Finally, BRDs in SMARCA2/4 (family VIII) (SWI/SNF related, Matrix associated, Actin dependent Regulator of Chromatin) and the scaffolding protein polybromo (bromodomain domain 5) have been selectively targeted by PFI-3^33^.

The chemical probe set presented here comprises 25 tool compounds covering 29 (50%) of all human BRD proteins (**Supplemental Table S1**). Each chemical probe interferes with the binding of its respective BRD(s) acetyl-lysine target sequences including the major acetylation sites described as BRD binding sites on histone proteins (**Figure 2**)^6^.

**Figure 2:**
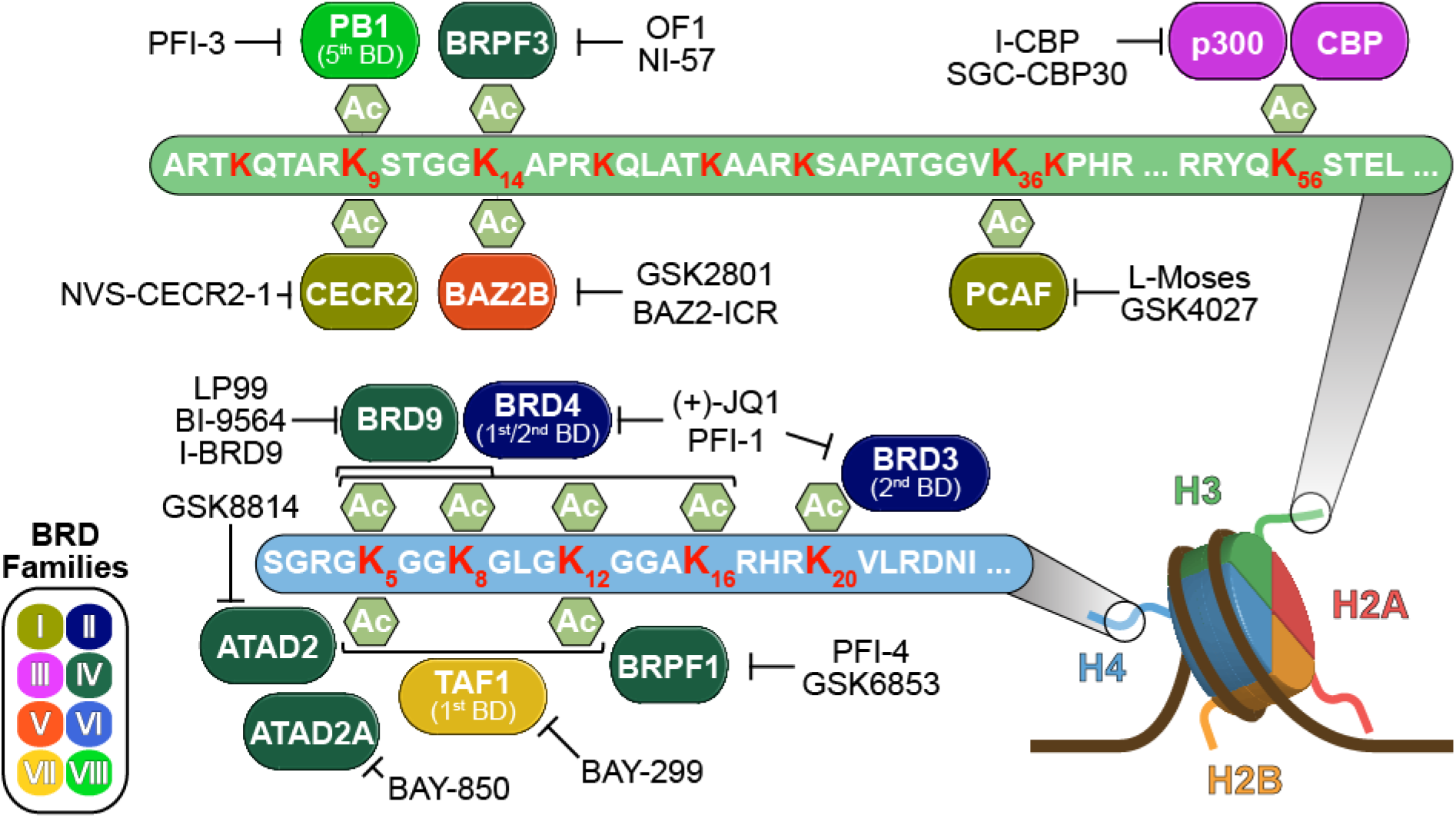
Schematic representation of bromodomain mediated interaction on histone tails that can be inhibited by the current set of chemical probes. Major sites of lysine acetylation histone H3 and H4 are highlighted in red.

### Selectivity of bromodomain chemical probes

The selectivity of chemical probes targeting diverse BRDs has been previously evaluated against a comprehensive set of recombinant human BRDs employing a temperature shift assay. This assay format makes use of the increase in the melting temperature (ΔT_m_) of a protein domain when complexed with a potent ligand^56^. However, intrinsic stability and other properties of proteins influence the magnitude of the observed temperature shift. Using bromosporine (BSP), a promiscuous BRD inhibitor^32^ we evaluated selectivity screens against a panel of BRDs employing the BROMOscan^®^ ligand binding assay, as well as isothermal titration calorimetry (ITC) and thermal melt assays (**Figure 3**). We used ITC as a standard for the accurate determination of binding constants, given its capacity to directly measure ligand binding in solution. All three assays resulted in comparable data (**Figure 3B**). However, while correlation between ITC and BROMOscan^®^ data was excellent (**Figure 3D**), some BRDs exhibited smaller than expected T_m_ shifts based on their binding constants determined by ITC (**Figure 3E**). In particular, BSP showed only modest T_m_ shifts against TAF1L(2) and BRD9 and a relatively large shift against CBP, compared to the directly determined ITC dissociating constants (K_D_s). Encouraged by the accuracy of the BROMOscan^®^ assay, we screened 15 chemical BRD probes against all 42 BRD-containing proteins and determined a total of 626 dose response curves (Supplemental Table S2). In addition to the BRD-probe set, we included three closely related variants of chemical probes within our set, CBP30-298 and CBP30-383 which are closely related to SGC-CBP30, as well as PFI-3 D1, a close derivative of PFI-3 (**Supplemental Figure S1**)^29,33,57^. However, while CBP30-related BET off-target effects were also apparent in the two additional CBP30 derivatives, the exclusive selectivity of PFI-3 towards SMARCA2/4 and PB1 was maintained in the derivative PFI-3 D1. Interestingly, the Kac mimetic salicylic acid head group of PFI-3 and its derivatives showed selectivity for this bromodomain subfamily. This striking observation has been rationalized by the unique binding mode of family VIII inhibitors that penetrated deeper into the Kac binding site leading to displacement of water molecules that are maintained in other BRD-inhibitor complexes^58, 59^. In summary, BROMOscan^®^ offers a robust platform for accurate K_D_ determination of BRD inhibitors, and chemical probes screened here maintained at least a 30-fold selectivity window against BRD in other families.

**Figure 3:**
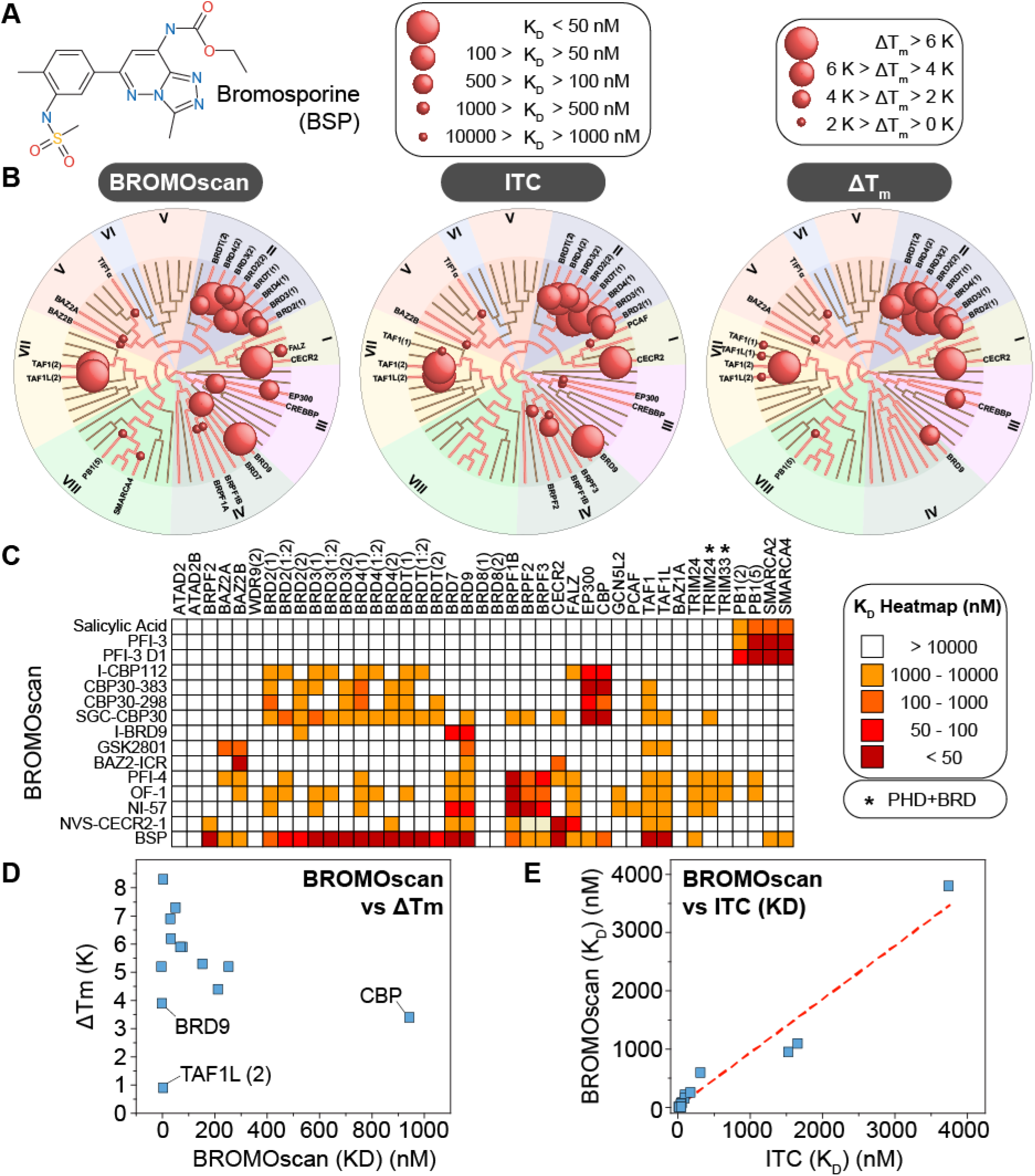
Selectivity of bromodomain chemical probes and assay comparison. **A)** Structure of bromosporine (BSP). **B)** Structural/phylogenetic dendrograms quantifying binding affinities of BSP to human BRDs measured by BROMOscan^®^ (left), ITC (middle) and Tm assays (right). Affinities and Tm shifts are mapped to the phylogenetic tree using sphere of variable size as indicated in the inset. Screened targets are annotated on the dedrograms. **C)** Heatmap of measured BROMOscan^®^ K_D_ values. **D)** Correlation of dissociation constants (K_D_) measured by ITC and BROMOscan^®^. **E)** Correlation of T_m_ shifts and dissociation constants (K_D_) measured by BROMOscan^®^.

### BET chemical probes block muscle cell differentiation by down-regulation of myogenin

Several BRD-containing proteins are known to be essential for different types of cellular differentiation, including myogenesis^33,34,57,60^. During myogenesis, differentiating myoblasts fuse to form multinucleated myotubes/myofibers, a process that plays an important role in development and regeneration, and BET proteins have recently been implicated in this process^61^. Given the importance of muscle regeneration in human health, we were interested in determining whether BRD inhibition would modulate the ability of muscle progenitor cells to undergo terminal differentiation. To explore this, the muscle progenitor C2C12 cell line was cultured in conditions of low serum to initiate differentiation in the presence of diverse BRD inhibitors from the chemical probe-set described here. Strong inhibitory effects of myoblast differentiation were seen for the BET inhibitor JQ1 as well as the pan-BRD inhibitor BSP, but not for other BRD inhibitors (**Figure 4A and B**). To gain insight into the transcriptional effects resulting from BRD inhibition, we performed gene expression analysis (**Figure 4C**). In agreement with our observations on myoblast differentiation, treatment with JQ1 and BSP resulted in significant changes in gene expression levels. Importantly, promiscuous targeting of BRD-proteins using BSP resulted in almost identical changes in gene expression compared to JQ1, strongly suggesting that the inhibition of differentiation is due to BET BRDs and not due to inhibition of other BRD-containing proteins. In agreement with this observation, expression changes resulting from targeting of other specific BRD-inhibitors outside the BET family were negligible (**Figure 4C**). Most significantly, differential expression analysis identified several anti-proliferative and anti-inflammatory genes downregulated including proteins modulating interferon response such as IFIT3 (interferon induced protein with tetratricopeptide repeats 3), interferon induced GTP hydrolases (GBPs) and USP18 (Ubiquitin Specific Peptidase 18) (**Figure 5A and B**). Gene-set enrichment analysis (GSEA) resulted in strong signatures for anti-inflammatory pathways and cell cycle regulators as well as myogenesis (**Figure 5C; Supplemental Figure S2**). Gene Ontology (GO) analysis corroborated these observations identifying enriched biological processes relating to cell cycle and mitosis as well as immune system processing and innate immune response (**Figure 5D, Supplemental Figure S2**). In particular, transcription of genes regulating expression of myosin light- and heavy-chains as well as regulators of myosin, such as myosin light chain kinase (MLCK) and myotonic dystrophy associated protein kinase (DMPK) and the tintin-associates proteins myomesin 1 and 2, was strongly suppressed. Key regulators of cellular fusion, important for the late stages of myogenesis, such as the protease ADAM12 were also affected. Importantly, we observed strong down-regulation of the muscle-specific basic-helix-loop-helix transcription factor myogenin (myogenic factor 4), a protein whose induction acts as a “point-of-no-return” in myogenesis by inducing cell cycle exit and activation of muscle-specific genes^62–65^. These observations suggest therefore that transient inhibition of BET bromodomain-containing proteins may be a means to delaying myoblasts from undergoing terminal differentiation. Interestingly, we did not observe transcriptional regulation of MYC and its target genes (**Supplemental Figure S2**), a gene expression response that is frequently used as a marker for BET inhibition in cancer highlighting the context dependent effect of BET inhibitors in different tissue types^18^.

**Figure 4:**
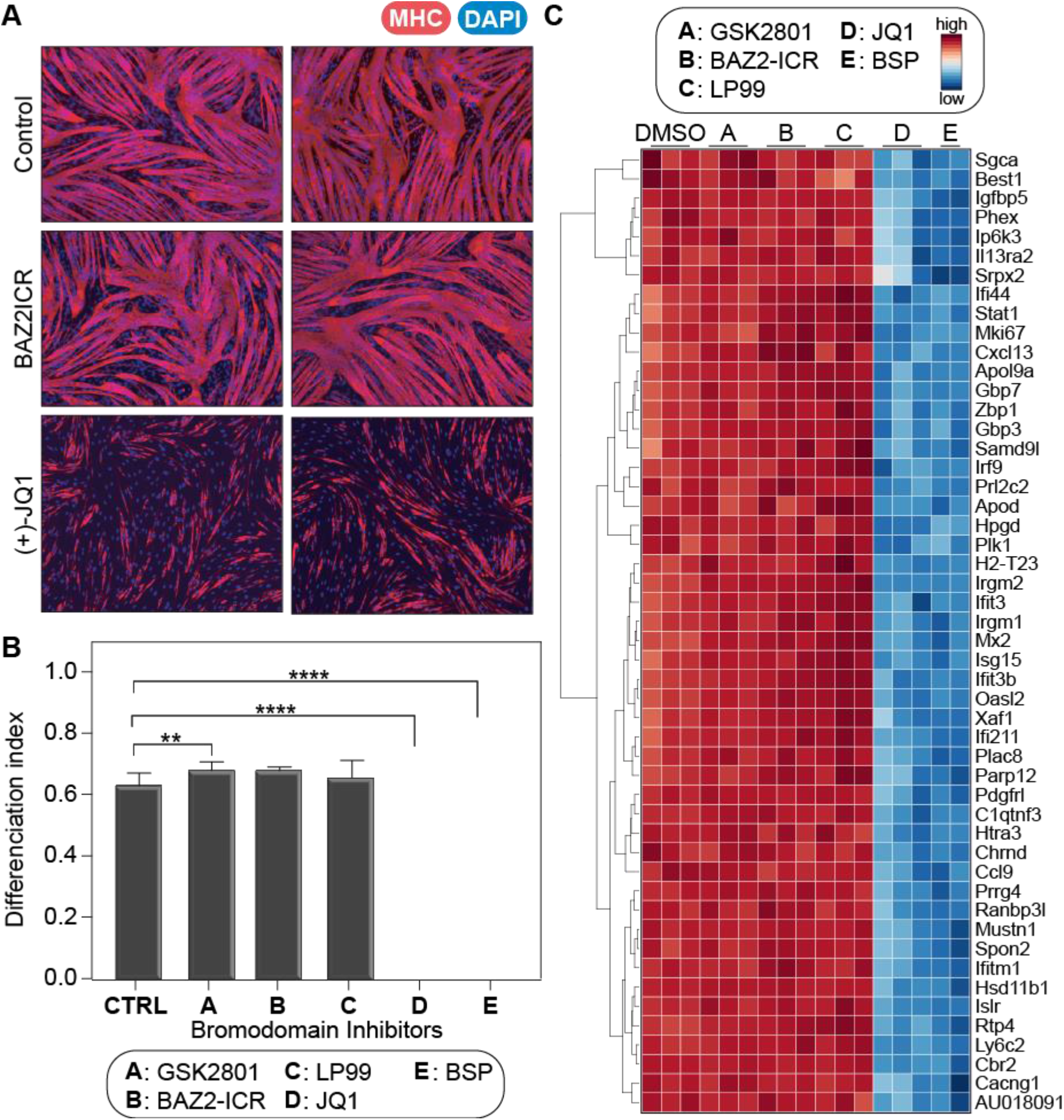
Influence of bromodomain inhibition on C2C12 myoblast differentiation. **A)** Fluorescent images of myotubes cultured in differentiation media (DMEM containing 2% horse serum, 10 μg/ml insulin and 10 μg/ml Transferrin) in the presence and absence of bromodomain chemical probes. Cells were allowed to differentiate for 48 hrs before they were processed for immunofluorescence staining with α-myosin heavy chain antibody (Red). Nuclei are stained blue with DAPI. **B)** Quantitation of differentiated cells after inhibitor treatment. Post-immunofluorescence, differentiation index was calculated by counting the number of nuclei in myosin heavy chain expressing myotubes divided by the total number of nuclei per field. **C)** Heatmap of the top 50 statistically significant genes that were differentially expressed (using Benjamini-Hochberg adjusted P-value < 0.001) following 12 h treatment with specific BRD inhibitors.

**Figure 5:**
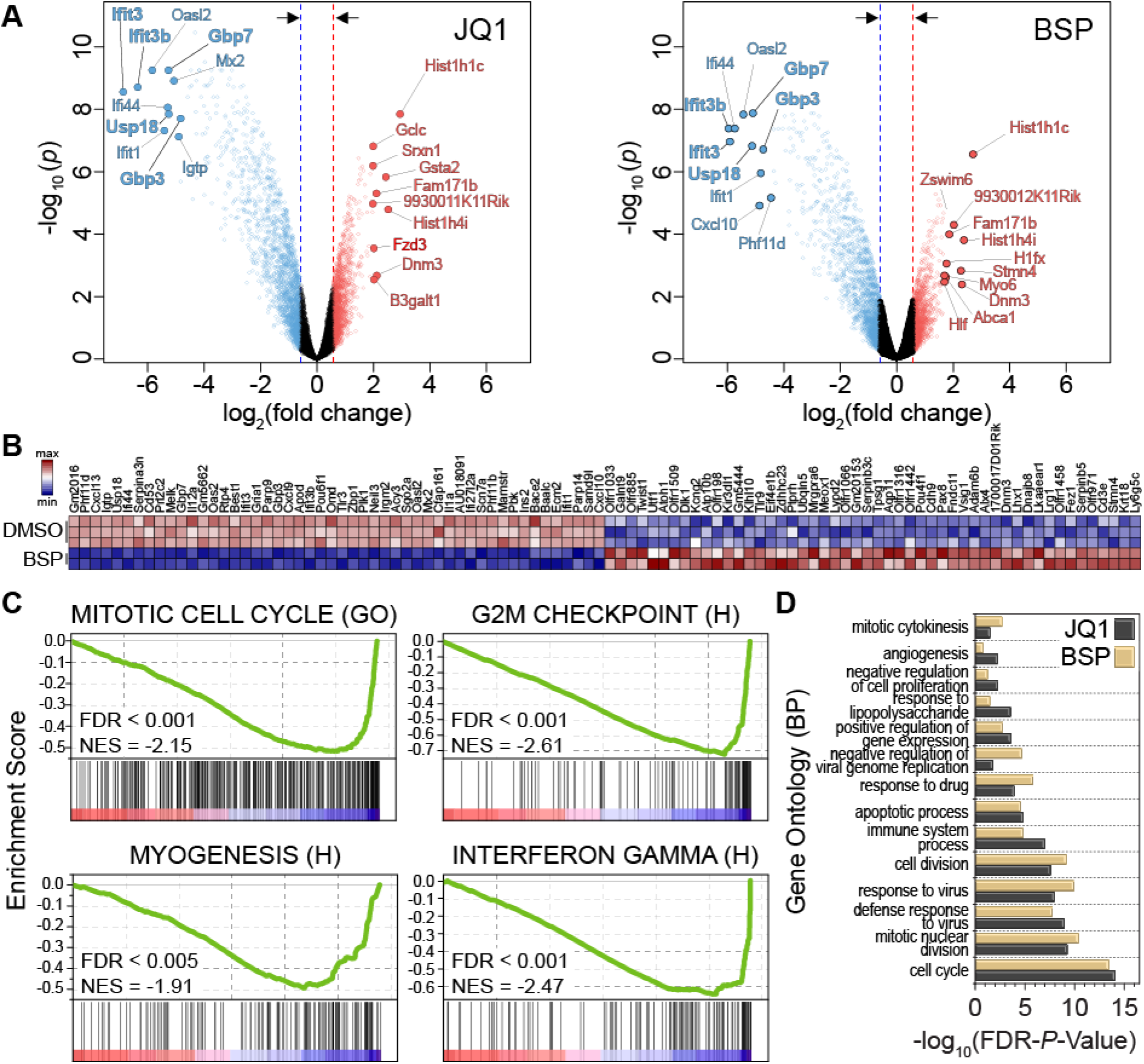
Transcriptional response to BSP and JQ1 in C2C12 myoblasts. **A)** Volcano plot of differentially expressed genes following 24 h treatment with BSP (left) or JQ1 (right) in C2C12 myoblasts. The top 10 genes are sorted by their fold-change and are highlighted and colored in red (up-regulated) or blue (down-regulated). **B)** Heatmap of the top 50 up/down regulated genes in C2C12 myoblasts following 12 h BSP treatment based on 2-sided signal to noise ratio (SNR) score and *P* < 0.05. Dark blue indicates lowest expression; dark red indicates highest expression, with intermediate values represented by lighter shades, as indicated in the inset. Data are column-normalized. **C)** GSEA demonstrating strong association with mitotic cell cycle (from the Gene Ontology MSigDB set, top left), G2M checkpoint (from hallmark MSigDB signatures, top-right), myogenesis (from hallmark MSigDB signatures, bottom-left) and interferon gamma (from hallmark MSigDB signatures, bottom-right) down-regulation signatures, following 12 h treatment of C2C12 cells with BSP. The plots show the running sum for the molecular signature database gene set within the C2C12/BSP data including the maximum enrichment score and the leading edge subset of enriched genes. Normalized Enrichment Scores (NES) and False Discovery Rates (FDR) are annotated in the insets. **D)** Gene Ontology (GO) enrichment (biological processes) for differentially expressed genes following 12 h treatment with JQ1 or BSP (calculated from differentially expressed genes with a Benjamini-Hochberg adjusted P-value < 0.001 and fold change > 1.5).

### BRD chemical probes reveal a metabolic/epigenetic circuit involving HBO1 in TNBC

Triple negative breast cancer (TNBC) accounts for almost 20% of breast malignancies and is characterized by lack of expression of the estrogen receptor (ER) and progesterone receptor (PR) and absence of HER2 amplification^66^. Due to the lack of targeted therapies, patients with TNBC have a poor survival rate and a larger likelihood of distant recurrence and death within five years of diagnosis^67–70^. Recent studies showed that BET inhibitors such as JQ1 are effective against TNBC by specifically downregulating genes required for tumor growth and progression^16, 71^. However, systematic investigations of the effects of other BRDis have not been evaluated in TNBC. To explore the potential anti-proliferation effects of BRDi, we profiled the viability of ten TNBC cell lines in the presence or absence of our BRD probe set. In agreement with previous studies, BET antagonists, including JQ1 and PFI-1, display strong antiproliferative effects on all the TNBC cell lines (**Figure 6A**), likely due to BETi effects on super enhancer-dependent transcription^15, 16^. However, no significant growth inhibitory effects were observed for the remaining BRDis.

**Figure 6:**
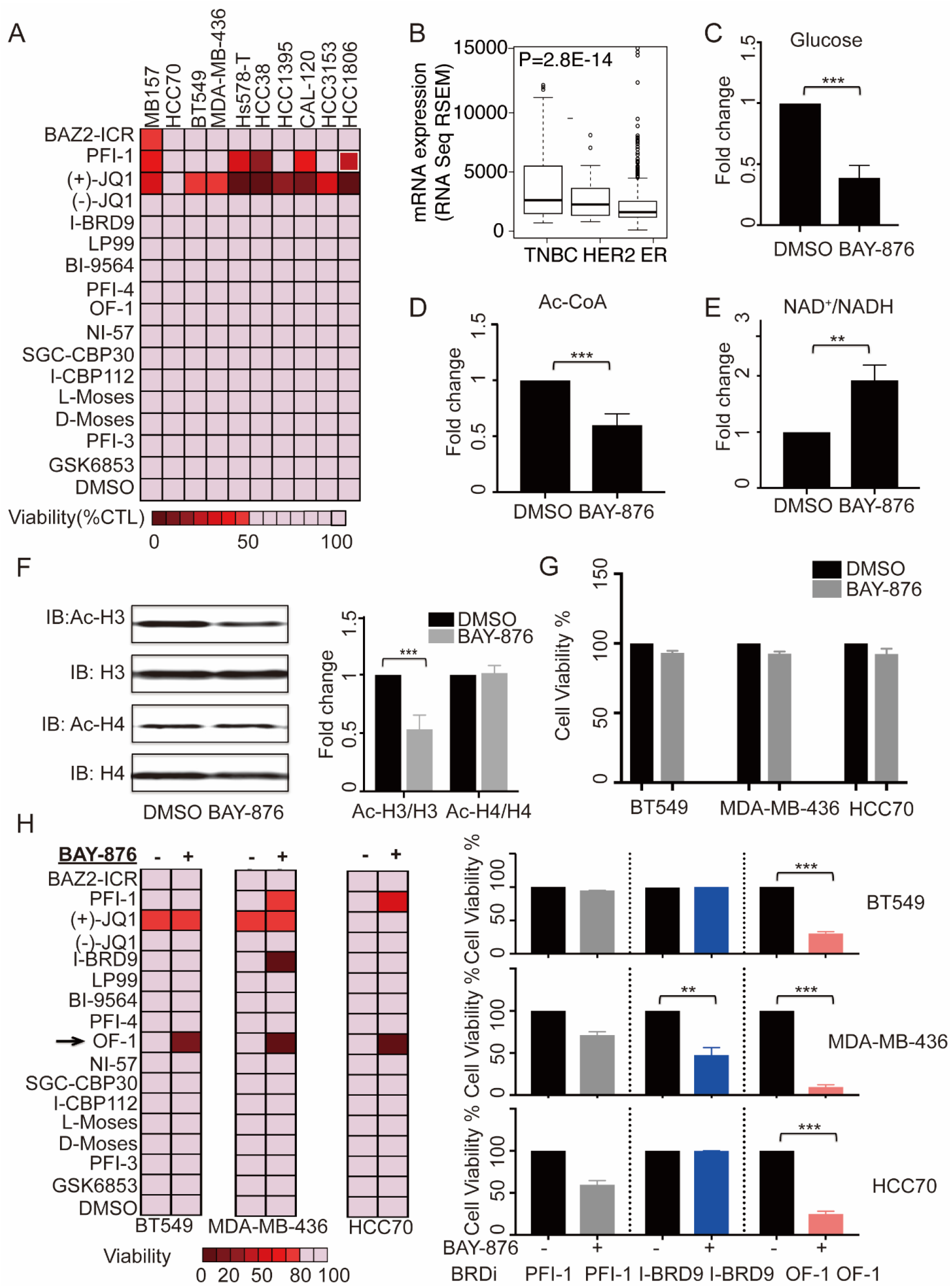
BRD inhibitors leverage metabolic adaptions induced by pharmacological inhibition of GLUT1 with BAY-876 in TNBC. **A**) BRD inhibitor screening across ten TNBC lines. Cells were treated with indicated BRD inhibitors at 3μM for 7 days. Confluency was measured using IncuCyte ZOOM live cell imaging device. **B)** GLUT1 (SLC2A1) mRNA expression in PAM50 based breast cancer subtypes. Gene expression data was downloaded from the publicly accessible TCGA (The Cancer Genome Atlas) portal. **C)** Glucose uptake in MDA-MB-436 cells in response to BAY-876 treatment relative to vehicle. MDA-MB-436 cells were treated with DMSO or 3μM BAY-876 for 5 days. Graph indicate mean and error bars denote standard deviation from three independent assays and p value computed using Benjamini-Hochberg t test; *** p<0.001 **D)** Effects of BAY-876 treatment on intracellular acetyl-CoA level. Graph indicate mean and error bars denote s.d. from three independent assays and p value computed using Benjamini-Hochberg t test; *** p<0.001 **E)** Effects of BAY-876 treatment on intracellular NAD+/NADH level. Graph indicate mean and error bars denote standard deviation from three independent assays and p value computed using Benjamini-Hochberg t test; ** p<0.01 F) Immunoblot analysis of H3 and H4 acetylation in MDA-MB-436 cells before and after BAY-876 treatment. p value computed using Benjamini-Hochberg t test; *** p<0.001. **G)** Cell viability effects of BAY-876 treatment on three representative cell lines. **H)** Combinatorial screening of BRD inhibitors with or without 3μM BAY-876 across ten TNBC lines. Cell viability was obtained from the endpoint Incucyte scanning (left). Heatmap of the combinatorial screening results; (Right) Cell viability after treatment with three potential BRDi candidates in the presence or absence of BAY-876 in three representative cell lines. Graph indicate mean and error bars denote standard deviation from three independent assays and p value computed using Benjamini-Hochberg t test; ** p<0.01; *** p<0.001.

Metabolism can directly impact cellular epigenetic landscapes and alter responses to chemo-therapeutics^72^. In particular, acetylation of histones relies on the availability of the universal acetyl donor metabolite acetyl-CoA, which is biosynthesized by breakdown of carbohydrates through the glycolytic pathway. Many TNBC cell lines display a classical Warburg metabolism with up-regulated glucose uptake to fuel their bioenergetics and biosynthetic demands^73, 74^. We confirmed that the set of 10 TNBC cell lines investigated in this study had a range of metabolically distinct states and variable global levels of histone acetylation (**Supplemental Figure S3A-C**). Furthermore, compared to other breast cancer subtypes, TNBC cell lines have a higher glycolytic gene-expression signature, especially for glucose transporter I (GLUT1) expression (**Figure 6B**), and thus tend to be more sensitive to glucose depletion^75^. Accordingly, this led us to investigate whether disruption of glycolysis in TNBC lines can give rise to epigenetic vulnerabilities to BRDis.

Using the selective GLUT1 inhibitor, BAY-876^76^, we first confirmed that exposure to BAY-876 inhibits glucose uptake in TNBC lines (**Figure 6C, S3D**). We then assessed the impact of BAY-876 treatment on levels of metabolites related to glycolysis and relevant to histone acetylation level. Acetyl-CoA is at the crossroads of glycolysis and TCA cycles and is the cofactor for histone acetyltransferases (HATs)^77, 78^. Upon treatment with BAY-876, a decrease in absolute acetyl-CoA level was observed, suggesting that HAT activity might be perturbed by BAY-876 treatment (**Figure 6D**). Another essential player involved in metabolism and acetylation is the NAD+/NADH ratio, which is closely associated with energy status in cell and is thought to positively regulate the activity of sirtuins^79^. We observed an increase in the NAD+/NADH ratio which may lead to increased sirtuin activity, possibly also contributing to histone hypo-acetylation (**Figure 6E**). Having observed a change in metabolites known to be involved in protein acetylation in response to BAY-876, we next investigated whether there are corresponding changes in histone acetylation levels. Interestingly, we detected a reduction in the global levels of acetylated histone H3 (ac-H3), but not in the global levels of acetylated histone H4 (ac-H4), in response to BAY-876 treatment (**Figure 6F**). These results demonstrate that manipulating metabolic flux by inhibiting glucose uptake can specifically impact the acetylation on individual histones.

We next assessed whether altered histone acetylation induced by BAY-876 treatment could induce sensitivity to BRDi in TNBC cell lines. We performed a combinatorial screen on three TNBC lines with distinct glycolytic rates (**Supplemental figure S3 C**). BAY-876 treatment alone had no effect on these three TNBC lines (**Figure 6G**), but in combination with the chemical probes PFI-1, OF-1 and I-BRD9 we observed a significant decrease in viability (**Figure 6H**). The activity of OF-1 was of particular interest due to the strong synergistic effect across all three cell lines, whereas JQ1 and I-BRD9 combinations with BAY-876 showed efficacy in only one of the three tested TNBC lines (**Figure 6H**).

To better understand the mechanism of the BAY-876/OF-1 combination, we determined the IC_50_ values for OF-1 treatment in all three cell lines. In the presence of 3 μM BAY-876, we observed IC_50_ values for OF-1 in the range of 0.3-2 μM (**Figure 7A**). An increase of OF-1 sensitivity was also observed in response to glucose deprivation mimicking the effect of BAY-876 treatment (**Figure 7B**). The combinatorial effect of BAY-876 and OF-1 was more than additive because the BAY-876 concentration we used in this assay has no overt effect on cell growth based on colony formation assay (**Supplemental figure S3E**). We next examined whether the observed synergy is due to the induction of apoptosis by the combination. Indeed, apoptosis markers such as caspase 3/7 were induced at a significantly higher level in combination-treated cells compared with cells treated with either OF-1 or BAY-876 alone (**Supplemental figure S3F**). Thus, we conclude that OF-1 and BAY-876 are synergistic in suppressing the growth of TNBCs by inducing apoptotic cell death.

**Figure 7:**
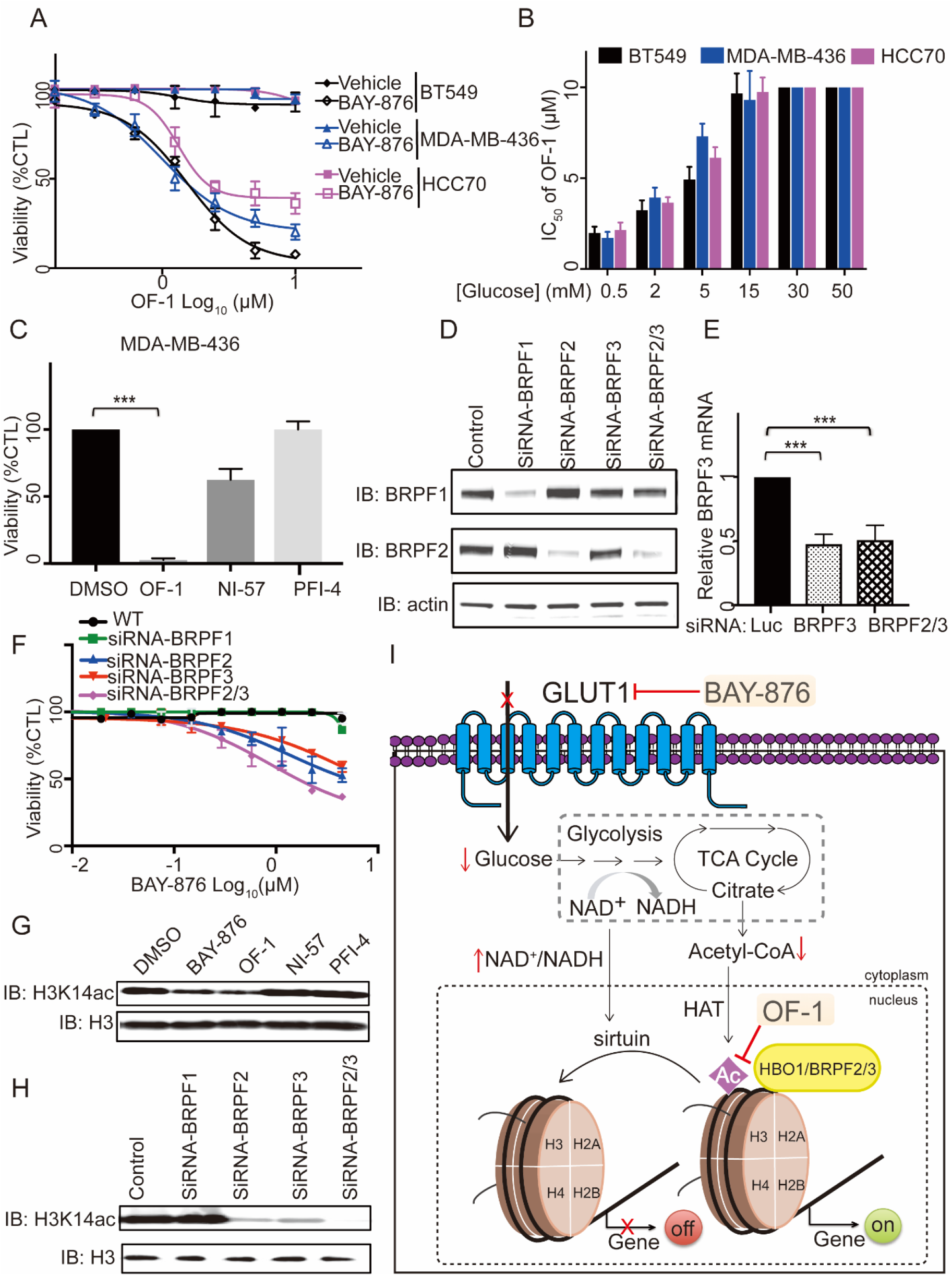
BRD chemical probes reveal a metabolic/epigenetic circuit involving HBO1 in TNBC. **A)** Dose-dependent curves for cells lines treated with indicated concentrations of OF-1 with or without 3μM BAY-876 for 7days. Graph indicate mean and error bars denote standard deviation from eight wells (from two independent assays. **B)** IC_50_ values of OF-1 in cells cultured under a range of glucose concentrations. **C)** Viability of MDA-MB-436 cells treated with indicated 3μM BRPF inhibitors: OF-1, NI-57 and PFI-4 for 7 days. Graph indicate mean and error bars denote standard deviation from eight wells (from two independent assays) and p value computed using Benjamini-Hochberg t test; *** p<0.001. **D)** Immunoblot validation of BRPF knockdown in MDA-MB-436 cells. (E) RT-qPCR validation of BRPF knockdown in MDA-MB-436 cells. p value computed using Benjamini-Hochberg t test; *** p<0.001. **F)** Dose-dependent response of BAY-876 in BRPF knockdown cell lines. **G)** Immunoblot analysis of H3K14 acetylation in MDA-MB-436 cells following BAY-876 and BRPF inhibitor treatment for 5 days. **H)** Immunoblot analysis of H3K14 acetylation in MDA-MB-436 knockdown lines. I) Schematic illustration of a metabolic/epigenetic circuit involving glucose transporter 1 and HBO1.

OF-1 inhibits Kac binding of the BRDs of BRPF1, BRPF2 and BRPF3. In order to deconvolute which BRD proteins are responsible for the observed phenotype with BAY-876, we compared the combinational effect on TNBC cell viability with another potent pan BRPF inhibitor NI-57, and the selective BRPF1 inhibitor PFI-4. Co-treatment with BAY-876 and OF-1 led to the strongest reduction on the cell viability, whereas other combinations were less effective (**Figure 7C**). Moreover, no significant effect was observed in the PFI-4 combination, which excludes the role of BRPF1 in the synergy effect. In agreement with these results, siRNA knockdown of BRPF1 did not change the cell sensitivity to BAY-876 treatment (**Figure 7D-F**). Notably, compared to the single BRPF2 or BRPF3 knockdown, the cells with dual BRPF2/3 knockdown are more sensitive to BAY-876 treatment (**Figure 7F**). Together, these results demonstrate that the observed synergy effects are due to the inhibition of BRPF2 or/and BRFP3 by OF-1.

BRPF2 and BRPF3 are components of the HBO1 (KAT7) acetyltransferase complex, while BRPF1 preferentially participates in the MOZ/MOF complex^80, 81^. Furthermore, BRPF2/3/HBO1 complexes were also shown to be important and specific for the HAT activity toward H3K14, whereas the BRPF1 complexes have high specificity on H4 acetylation marks^81^. To explore whether the inhibition of BRPF2 or/and BRPF3 has any effect on histone acetylation, we measured the H3K14ac level upon treatment with BRPF BRD chemical probes, or in BRPF knockdown cell lines. Compared to NI-57 and PFI-4, the acetylation of H3K14 was significantly decreased in the presence of OF-1 (**Figure 7G, S2G**). Likewise, dual BRPF2/3 knockdown displayed the strongest reduction of H3K14ac compared to BRPF2 or BRPF3 knockdown alone (**Figure 7H, S2H**). Taken together, these data are consistent with a model in which inhibition of glucose up-take by BAY-876 and antagonism of HBO1 subunits BRPF2/3 by OF-1, both converge on the same histone marks, leading to synergistic cross talk between metabolic and epigenetic pathways (**Figure 7I**).

## Discussion

Recent effort by our laboratories and others established a comprehensive set of epigenetic probe molecules for selective targeting of acetyl-lysine dependent reader domains. We believe that this is significant achievement considering that apart from a number of fragment-like small molecules, no BRD had been targeted before the first potent BET inhibitors were disclosed in 2010^8, 9^. In particular, BET inhibitors had a major impact on basic and translation research as demonstrated by the large number of research papers that have been published using these reagents and more than 20 clinical trials that are currently registered^10^. However, BRD inhibitors outside the BET family have not been validated as potential drug targets. Here we provide data showing that inhibition of BRPF BRDs in combination with selective inhibitors of glucose transport might be beneficial for the treatment of TNBCs. Earlier studies demonstrated also synergies of BRD inhibitors with other drugs, such as CBP/p300 inhibitors acting synergistically with BET inhibitors as well as cytotoxic agents and dexamethasone in leukemia^34^. In addition, BET inhibitors showed synergy in cancer models in combination with HDAC inhibitors^82–84^ as well as kinase and PARP inhibitors^85–87^. The combination of different inhibitors might also be important in overcoming drug resistance which has been observed in cells treated with BET inhibitors^13^. The reported surprising dual activity of kinase and BET inhibitors suggests that potent activity for both bromodomain and kinases could be designed into a single inhibitor^88–91^.

The profiling data provided here offers a comparison of inhibitor potency and selectivity across the BRD family. We found good correlation of BROMOscan^®^ assay data with K_D_s determined in solution by ITC, whereas the magnitude of T_m_ shits across the BRD family may vary depending on the intrinsic stability of each BRD. However, as an analytical tool the Tm shift assay provides a good platform for assessment of selectivity when hits are carefully followed up using orthogonal binding assays. Some probe compounds were exclusively selective while others, such as the CBP/p300 probes I-CBP112 and CBP30, showed significant BET activity (Supplemental Table S2). Thus, care should be taken when these probes are used in cellular assays. We recommend that probe concentration not higher than 3 μM for I-CBP112 and 2.5 μM for CBP30 are used in cell-based assays and that BET inhibitors are included as controls.

Even though the coverage of the bromodomain family with chemical probes is now quite good, there are still a number of BRDs for which no selective or even non-selective inhibitors are available. Many of these BRDs have unusual Kac binding sites, for instance in some BRDs the conserved Asn is replaced by Ser, Thr and Tyr residues^6, 92^. No specific Kac containing sequences have been reported binding to these BRDs limiting the development of Kac competitive assays. Some of these BRDs may also not recognize Kac-containing sequences at all. Other BRDs have less druggable binding sites making the development of high affinity chemical probes challenging. There are now also structurally diverse Kac-binding domains called YEATs domains that have not been targeted^93^. Some of these domains recognize also crotonylated lysine residues in addition to Kac containing sequences^94^. It is therefore likely that the arsenal of chemical probes for these reader domains will continue to grow in the future.

Most bromodomain containing targets are complex multi-subunit containing molecules which also contain histone- and chromatin-interacting proteins. For some BRD containing proteins, such as BET proteins, chemical antagonism of Kac binding is sufficient to displace the target protein from its intended chromatin loci^15^. In other cases, such as for p300/CBP and SMARCA2/4 containing complexes, it appears that BRD antagonism is insufficient to displace the entire complex from chromatin^33, 34^. Thus, BRDi targeting complex chromatin proteins are not likely to replicate genetic knock down studies deleting the full-length protein^33, 95^. We believe that this chemical probe tool set will be an excellent resource for understanding the role of specifically targeted BRDs within larger chromatin complexes and may unravel novel opportunities for translational research projects.

## Acknowledgements

The authors are grateful for support by the SGC, a registered charity (number 1097737) that receives funds from AbbVie, Bayer Pharma AG, Boehringer Ingelheim, Canada Foundation for Innovation, Eshelman Institute for Innovation, Genome Canada, Innovative Medicines Initiative (EU/EFPIA) [ULTRA-DD grant no. 115766], Janssen, Merck KGaA Darmstadt Germany, MSD, Novartis Pharma AG, Ontario Ministry of Economic Development and Innovation, Pfizer, São Paulo Research Foundation-FAPESP, Takeda, and Wellcome [106169/ZZ14/Z] and the DFG funded centre of excellence (CEF) at Frankfurt University. We thank the High-Throughput Genomics Group at the Wellcome Trust Centre for Human Genetics (funded by Wellcome Trust grant reference 090532/Z/09/Z) for the generation of Gene Expression data and the Medical Research Council (MRC grant MR/N010051/1 to PF) for funding.

